# Resting state fMRI based target selection for personalized rTMS: stimulation over the left DLPFC temporarily alters the default mode network in healthy subjects

**DOI:** 10.1101/344390

**Authors:** Aditya Singh, Tracy Erwin-Grabner, Grant Sutcliffe, Andrea Antal, Walter Paulus, Roberto Goya-Maldonado

**Affiliations:** Systems Neuroscience and Imaging in Psychiatry, Department of Psychiatry and Psychotherapy of the University Medical Center Göttingen; department of Clinical Neurophysiology of the University Medical Center Göttingen

## Abstract

High frequency repetitive TMS (HF-rTMS) to the left dorsolateral prefrontal cortex (DLPFC) is an approved option for the treatment of depression, but there is also much variability in the resulting antidepressant response. This variability is believed to arise from various factors, an important one of which being the method by which rTMS is targeted to the optimal stimulation site in the left DLPFC. To more accurately target HF-rTMS at left DLPFC, we present a new method for target selection based on individual RS-fMRI data. We show in 23 healthy subjects that the new proposed method of target selection is reproducible and yields left DLPFC targets whose functional connectivity correlates more negatively with subgenual anterior cingulate cortex (sgACC) than targets based on standard MNI coordinates. Since previous work has highlighted higher negative connectivity with the sgACC as an important feature of targets for higher antidepressant effect of HF-rTMS, the targets selected by the new method can be expected to lead to a higher therapeutic response. Additionally, the mechanism of action of an entire single session of HF-rTMS (3000 pulses) in healthy subjects has not been reported. We show significant decreases in functional connectivity of the default mode network in sgACC and ventral striatum (vStr) regions, peaking at 27-32 minutes after stimulation. Also, we report a negative correlation between the magnitude of this decrease in the right sgACC and the harm avoidance domain measure from the Temperament and Character Inventory (TCI). This finding points towards the possibility of using the harm avoidance measure as a predictor of HF-rTMS response. In addition, the decreased functional connectivity of the default mode network in right nucleus accumbens (NAcc) correlates with a short-term decrease in self-rated negative emotions from the Positive and Negative Affect Schedule (PANAS) i.e. the lower the functional connectivity of right NAcc with the default mode network, the lower the reported perception of negative mood by the subjects. This suggests a mechanism by which changes induced by rTMS influence the perception of negative mood in recipients.

## Introduction

Transcranial Magnetic Stimulation (TMS) is a non-invasive tool that allows targeted stimulation of brain areas. TMS was first proposed as a brain stimulation tool by Barker and colleagues in 1985 (Barker et al., 1985) and functions according to Faraday’s principle of magnetic induction, whereby a rapidly changing electric current gives rise to a change in local magnetic field, which in turn induces an electrical current in the neurons, thus promoting or inhibiting spiking activity (Wagner et al., 2007). Stimulation is delivered in short ‘pulses’ and various pulse protocols are available, such as single pulses, paired pulses and repetitive pulses. Single and repetitive TMS (rTMS) protocols have been employed for a variety of clinical and non-clinical purposes, such as probing motor cortex excitability (Kujirai et al., 1993; Chen et al., 1997) and cognitive processes (Fadiga et al. 1995), treating chronic pain (Lefaucheur et al., 2001), movement disorder in Parkinson’s disease (Hamada et al. 2008), stroke (Mansur et al., 2015), and psychiatric disorders such as depression (Pascual-Leone et al., 1996), anxiety disorders (Cohen et al., 2004), obsessive compulsive disorder (Greenberg et al., 1997), schizophrenia (Hoffman et al., 2000), substance abuse (Mishra et al., 2010). Detailed evidence-based guidelines for the therapeutic use of TMS have been published elsewhere (Lefaucheur et al., 2014; Rossi et al., 2009).

In 2008, the FDA approved rTMS for the treatment of drug resistant depression after large scale studies confirmed efficacy in relieving depressive symptoms and improving patients’ quality of life (O’Reardon et al., 2007; George et al., 2010, Solvason et al., 2014). Over time, more studies have explored the therapeutic benefits of other forms of rTMS, such as intermittent theta-burst stimulation (Cheng et al., 2016; Li et al., 2014; Blumberger et al., 2018; Bulteau et al., 2017; Duprat et al., 2016). Despite the proven benefits of rTMS, there remains large variability in the interindividual antidepressant effects of rTMS treatments, which is hypothesised to stem from differences in age (Peinemann et al., 2001), white matter connectivity (Quentin et al., 2013), genetic polymorphisms (Kleim et al., 2006; Cheeran et al., 2008) and the method by which rTMS is targeted (Ahdab et al., 2010; Herwig et al, 2001; Herwig et al., 2003; Rusjan et al., 2010). Fitzgerald and colleagues (Fitzgerald et al., 2009) investigated methods of targeting and stimulating the DLPFC and concluded that targets defined on the 10-20 electroencephalographic (EEG) system or selected with the use of individual structural MRI images and applied with a neuronavigation system is better than those selected with the “standard procedure” of using the scalp location 5 cm anterior to the motor cortex (as described in Lefaucheur et al., 2014). Using data from functional MRI scans of individual subjects (Fox et al. 2012 and Weigand et al., 2017) have showed that therapeutic efficacy of targets for high frequency (HF) rTMS in depression is predicted by negative functional connectivity of the targets with the subgenual anterior cingulate cortex (sgACC). These studies along with others (Lefaucheur et al., 2007; Lefaucheur et al., 2010; Ruohonen and Karthu, 2010) imply that using brain imaging to guide targeting would be an effective strategy in increasing the efficacy and reliability of therapeutic rTMS.

We argue that for maximal therapeutic benefits it is important to incorporate functional connectivity features when selecting rTMS targets, and to guide stimulation accurately to these targets. In this study, we apply a novel method of resting state fMRI based target selection for personalized HF-rTMS in healthy volunteers. This new method employs the individual subject’s resting state fMRI (RS-fMRI) data at baseline to select the most promising spatial spot over the left dorsolateral prefrontal cortex (DLPFC), in order to indirectly target the subgenual anterior cingulate cortex (sgACC), which is of major therapeutic significance in depression (Fox et al., 2012, Fox et al., 2013, Weigand et al., 2017).

Several studies have explored the brain connectivity changes post rTMS treatment in depression patients (Taylor et al., 2018; Liston et al., 2014) and numerous other studies have looked at the effects of rTMS on healthy subjects (Rahnev et al., 2013; Ji et al., 2017; Halko et al., 2014; Wang et al., 2014), although to our knowledge, no study to date has explored the mechanism of action of a complete 10 Hz rTMS session (about 40 min and with 3000 pulses delivered) in healthy subjects. An important recent study (Tik et al., 2017) explored the mechanism of 10 Hz rTMS in healthy subjects, but only applied about a third of the protocol (1200 pulses, 10 min) used clinically in depression. The fact that many of depression patients do not respond to rTMS treatment emphasizes the need to develop personalized approaches as well as understand the mechanism of action of a single full HF-rTMS session. Such insights could deliver novel insights and help open new pathways to boost the antidepressant effect of rTMS. This study addresses the mechanism of action of a complete 10 Hz rTMS session using RS-fMRI functional connectivity in a crossover, sham controlled study with healthy subjects. Taking into consideration the whole body of evidence indicating the importance of the default mode network in depression (Grecuis et al., 2007; Lit et al., 2013; Manoliu et al., 2013; Zhu et al., 2012, Van Tol et al., 2013; Sheline et al., 2010; Alexopoulous et al., 2012; Berman et al., 2011; Wu et al., 2011, Andreescu et al., 2013; Bluhm et al., 2009, Mulders et al., 2015, Liston et al., 2014) as well as the subgenual ACC in antidepressant responses, we hypothesized that functional connectivity changes would occur in this network. Moreover, our method individually targeted the functional relationships between the left DLPFC and the sgACC. Based on past research (Mulder et al., 1994; Ward et al., 2018; Quilty et al., 2017; Seon-Young et al., 2007), we investigated the possibility of predicting rTMS response using temperament and character inventory (TCI) trait parameters. We also expected temporary coupling changes in the sgACC to correspond to differences in self-rated emotional state (Abend et al., 2018; Drevets et al., 2008; Nitsche et al., 2012; Mondino et al., 2015: Leyman et al., 2009).

## Methods

### Participants

We recruited healthy male and female subjects between the ages of 18-65 with no current or prior psychiatric disorders evaluated by structured clinical interviews (see below). Exclusion criteria were current or history of neurological or psychiatric disorder, illegal drug use in the past month, current or past history of substance abuse or dependence, contraindications to the MRI scanner (e.g. metal parts in the body) or TMS application (e.g. epilepsy), pregnancy, history of traumatic brain injury, unwillingness to consent or to be informed of incidental findings, current use of anticonvulsant drugs, or prior TMS or ECT application in the past 8 weeks. The Ethics Committee of the University of Medical Center Goettingen approved the study protocol.

### Day 1

The subjects visited the lab on 3 occasions (Day1, Day2, and Day3). On Day 1 we provided a verbal and written description of the study. We obtained oral and written consent from all participants, and we anonymized all the information collected, including clinical interviews and psychological measures used to determine inclusion/exclusion in the study, demographic information and behavioural measures. Initial structural and RS-fMRI scans were recorded for electric field modelling and our novel method oc individualized target selection. During RS-fMRI, we instructed subjects to look at a fixation cross presented on a black background. RS-fMRI sessions were followed by an index finger tapping session (intermittent finger tapping when presented with a green or a red dot), data from which was used to determine the motor cortex location used for setting the motor threshold. Behavioural scales consisted of the Montgomery-Asberg Depression Rating Scale (MADRS), Beck Depression Inventory II (BDI II), Symptoms Checklist 90 (SCL-90), Young Mania Rating Scale (YMRS), Positive and Negative Syndrome Scale (PANSS), Barratt Impulsiveness Scale (BIS), Temperament and Character Inventory (TCI), Life Orientation Test-Revised (LOT-R), Mehrfachwahl-Wortschatz-lntelligenztest (MWT, vocabulary intelligence test) and handedness questionnaire (Oldfield, 1971).

### Target Selection

For real and sham rTMS on day 2 and day 3, we used the RS-fMRI scan from day 1 to identify a personalized target in the left dorsolateral prefrontal cortex (DLPFC) for each subject, using a novel selection process as detailed below.

Using SPM12 (http://www.fil.ion.ucl.ac.uk/spm/software/spm12/) and MATLAB (The MathWorks, Inc., Natick, MA, USA), we preprocessed the individual RS-fMRI data using standard steps: slice time correction, motion correction, individual gradient echo field map unwarping, normalization, and regression of white matter, cerebrospinal fluid and motion nuisance parameters (Figure 1a). We then temporally concatenated the data to perform group and individual independent component analyses with FSL 5.0.7 software (Jenkinson et al., 2012). The number of independent components was restricted to 17 based on the literature (Yeo et al., 2011). We identified the best fitting independent components (IC) covering: 1) the left DLPFC (IC-DLPFC) and 2) the sgACC (IC-ACC).

**Figure 1a:**
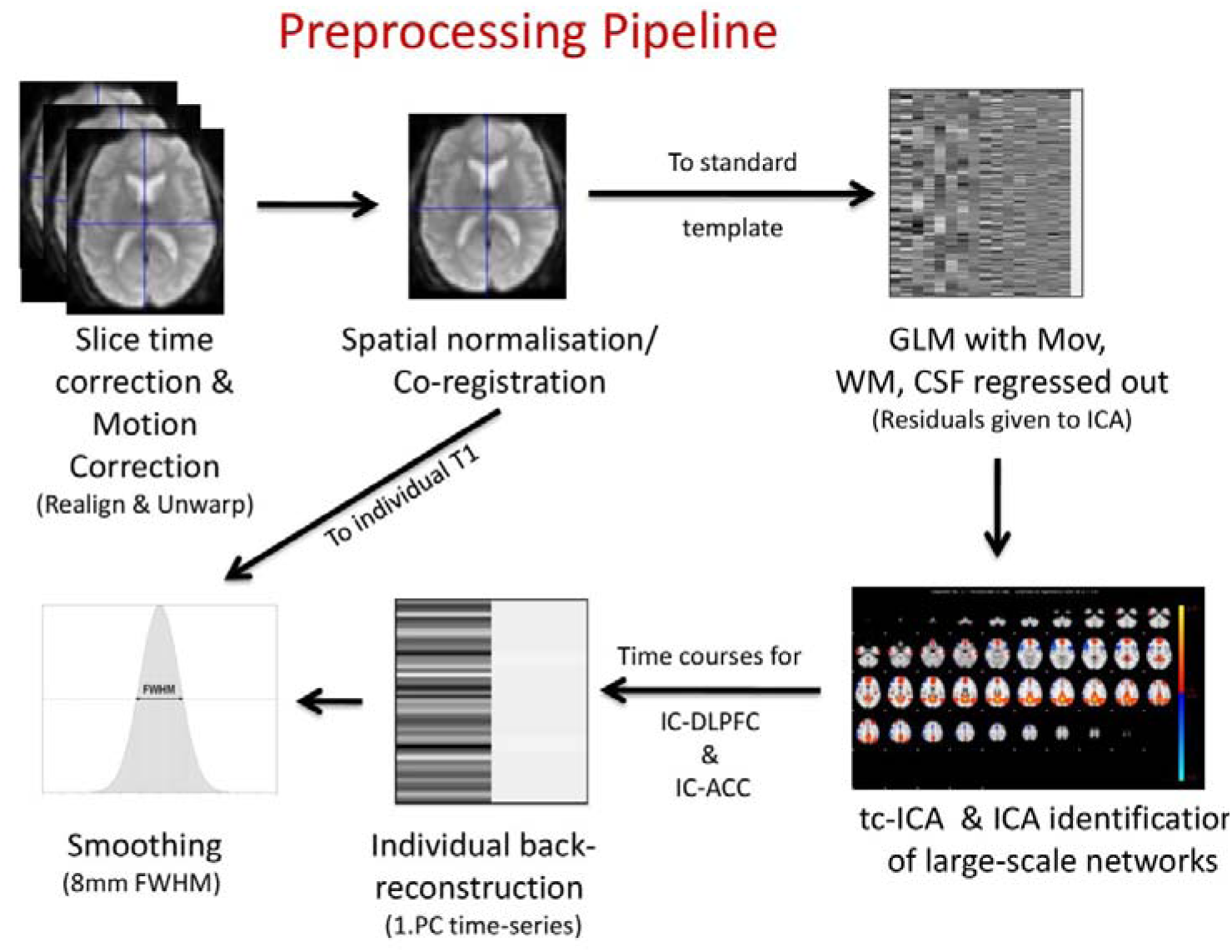
The two part figure demonstrates the target selection process. In the first part, the RS-fMRI data from the individual subject is pre-processed. We run ICA (both individually, with no. of components set to 17 and a group ICA, tc-ICA) on RS-fMRI data that is normalized. The relevant independent components (IC): IC-DLPFC and IC-ACC (see Methods section) are regressed in RS-fMRI data that is not normalized and is smoothened with an 8 mm FWHM kernel.

**Figure 1b:**
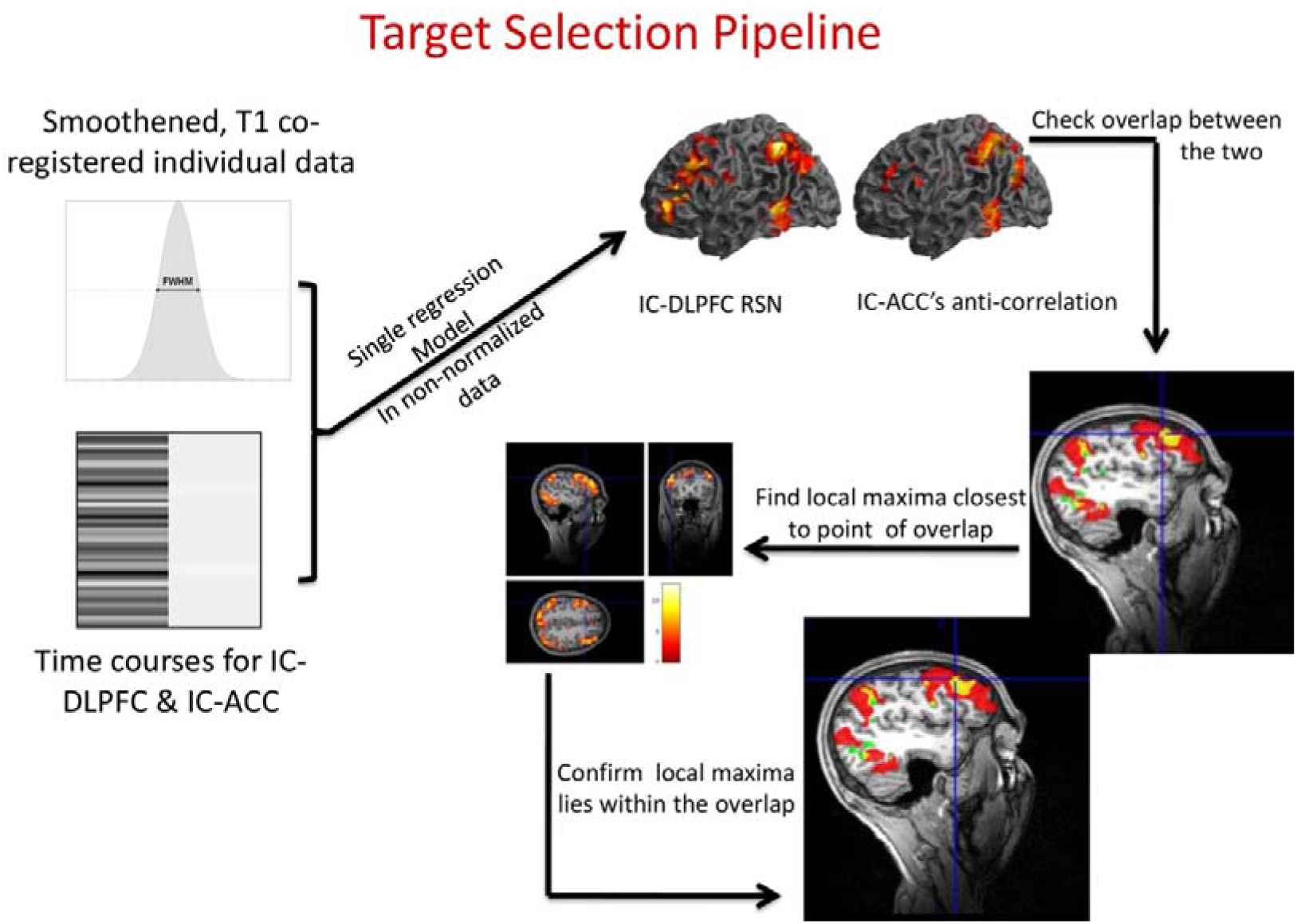
In the second part, the regressed IC in individualized data is overlapped using MRIcro and a point in the region of overlap within the left DLPFC is located. The maximal node closest to this point is identified in the IC-DLPFC using SPM12. This point becomes the target of stimulation, if it continues to lie within the overlap of IC-DLPFC and anti-correlation of IC-ACC.

**Figure 2:**
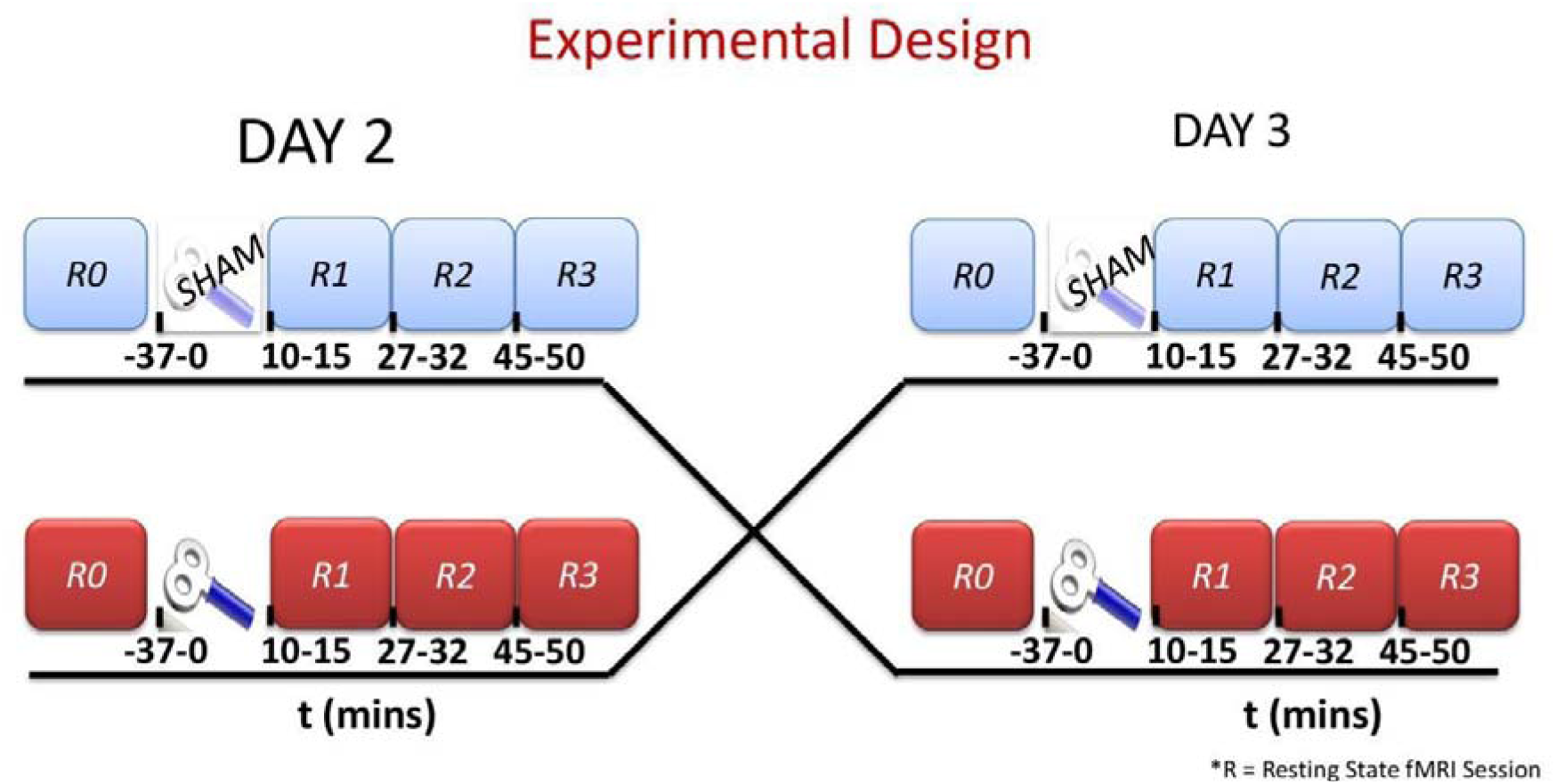
Experimental design of the study. Subjects underwent 4 RS-fMRI scans on each day. After the baseline RS-fMRI scan (R0), we delivered stimulation using 10 Hz rTMS. After the stimulation, there were 3 RS-fMRI scans: R1, R2, and R3 at 10 minutes, 27 minutes, and 45 minutes, respectively. For sham stimulation the MCF-B65 Cool coil was inverted by 180° and the non-stimulating side was used.

The ICs obtained were in standard MNI space. To reconstruct left DLPFC targets in individual brain spaces, we reran the preprocessing pipeline described above, but without normalizing the data and instead co-registering it to the subject’s T1 image, and additionally smoothing the data with a FWHM Gaussian kernel size of 8 mm. Next, we back reconstructed the IC-DLPFC and IC-ACC in the non-normalized RS-fMRI data to obtain these components in the individual anatomical space.

As previously reported in the literature (Fox et al., 2012 and Weigand et al., 2017), better clinical responses to rTMS in patients with depression are associated with higher negative functional connectivity between the sgACC and stimulation target in the left DLPFC. To incorporate this feature into our personalized target selection for rTMS, we overlaid individual negative correlation maps of the IC-ACC with the positive correlation maps of the IC-DLPFC (p<0.001) using MRIcron. The negative correlation of IC-ACC was thresholded leniently (p<0.01) to allow the detection of viable overlaps. We located a point within the overlap of these two maps and then identified the closest maximum nodes (strongest node) of the IC-DLPFC to this point in using SPM12. From the maximum nodes that lay in the overlap, the node with the peak connectivity within the IC-DLPFC was selected as the target for rTMS stimulation (Figure 1b). By doing this we incorporated two features that define an optimal target: the target is based on individual RS-fMRI data (Fox et al., 2013) rather than a group average and the target has a negative correlation to sgACC (Fox et al., 2012 and Weigand et al., 2017).

Also, note that even though we ran a temporally concatenated ICA on a group of RS-fMRI data that included the subject for which the target was sought, we used the ICs obtained from individual ICA results for target selection. We used the ICs from tc-ICA only in cases when individual ICA failed to yield any viable points, due to lack of an overlap. In the cases that we were unable to obtain an overlap using tc-ICA ICs, we repeated the day 1 RS-fMRI measurement and applied target selection to the new data. For cases in which this was necessary, we successfully identified targets when RS-fMRI measurement was repeated for all except one subject.

### Day 2 and Day 3

At the beginning of experiment on day 2 and day 3, we asked each subject to complete the Positive and Negative Affect Schedule (PANAS). Next, to determine individual resting motor thresholds (RMT) we stimulated the Abductor Pollicis Brevis (APB) by delivering single pulses of TMS at the primary motor cortex finger region identified during the day 1 finger tapping task, guided by an online neuronavigation system (Visor 1 software, ANT Neuro, Enschede, The Netherlands). We defined the RMT as the TMS stimulator output at which less than 5 out of 10 TMS single pulses resulted in an electromyography (EMG) response of more than 50 μV amplitude, and then set the stimulator to deliver 10 Hz rTMS at 110% of this threshold (see rTMS Stimulation section). We obtained a baseline RS-fMRI scan (R0), and then delivered HF rTMS stimulation (either real or sham in a counterbalanced way) to the subject at the previously selected target (see Target Selection section), guided by online neuronavigation. In the event of extreme scalp discomfort, the stimulation was stopped and the subject was excluded from further experiments.

Three additional RS-fMRI scans (R1, R2, R3) were obtained over a course of 50 minutes post HF rTMS to detect effects on brain resting state functional connectivity. The subjects again completed the PANAS immediately after leaving the MR scanner. This allowed us to document any short term changes in the self-rated emotional state.

After the end of scanning sessions and completion of the PANAS, subjects completed a poststimulation assessment which included the following behavioural scales and questionnaires: MADRS, BDI II, YMRS, PANSS, HAM-D. To assess the effectiveness of sham blinding, we also retrospectively, collected information about the perceived effects of condition on scalp sensation and mental state using a visual analog scale (VAS).

### rTMS Stimulation

We delivered 10 Hz rTMS using a MagVenture X100 with Mag-option and a MCF-B65 cooled butterfly coil. The stimulation parameters were: a train of pulses lasting 4 seconds followed by 26 seconds of rest period. We delivered 75 trains for a total of 3000 pulses in duration of 37.5 minutes (as described in O’Reardon et al., 2007). This is a full protocol session used in the clinic for the treatment of depression.

To deliver sham stimulation, we rotated the coil by a full 180° along the handle axis of the coil such that the stimulation side of the coil faced away from the scalp and the distance between the stimulation side and the scalp was larger than 5 cm. We also made measurements of the voltage induced on the “sham” side using a standard oscilloscope. The oscilloscope readings indicated that there was a very weak current strength produced by the sham side of the coil, and this negligible current did not elicit any motor responses, irrespective of how high the stimulator output was set to. Participants were blind to the control condition and simply received the information that we were testing two different stimulation protocols.

### Imaging Acquisition and Analysis

We acquired the structural (T1- and T2-weighted scans with 1-mm isotropic resolution) and functional data with a 3T MR scanner (Magnetom TRIO, Siemens Healthcare, Erlangen, Germany) using a 32-channel head coil. The gradient-echo EPI sequence had the following parameters: TR of 2.5 seconds, TE of 33 ms, 60 slices with a multiband factor of 3, FOV of 210 mm × 256 mm, 2×2×2 mm, with 10% gap between slices and anterior to posterior phase encoding. The RS-fMRI data was acquired with 125 volumes in approx. 5.5 minutes, whereas the finger tapping data was acquired in 103 volumes in approx. 4.5 minutes.

### RS-fMRI data analysis

Using SPM12 (http://www.fil.ion.ucl.ac.uk/spm/software/spm12/) and MATLAB (The MathWorks, Inc., Natick, MA, USA), we preprocessed the individual RS-fMRI data using standard steps: slice time correction, motion correction, individual gradient echo field map unwarping, normalization, and regression of white matter, cerebrospinal fluid and motion nuisance parameters. We then temporally concatenated the data to perform group independent component analysis with FSL 5.0.7 software (Jenkinson et al., 2012). We identified the best fitting independent component (IC) that resembled the default mode network. This IC was then back reconstructed in individual subjects’ normalized RS-fMRI data, r-to-z transformed and compared across the groups (Real [R0, R1, R2, R3] versus Sham [R0, R1, R2, R3]).

### Finger tapping fMRI data

We pre-processed the finger tapping data using slice time correction, motion correction, gradient echo field map based distortion correction, co-registration to the anatomical scan and smoothing with an 8 mm FWHM kernel. The onset times and durations for green dot (finger tapping) and red dot (rest period) were extracted from log files generated by Presentation software (Neurobehavioral Systems, Inc.) to create a block design of the experiment. Estimates of neural activity were computed with a general linear model (GLM) for each subject individually using SPM12. First-level contrasts were calculated for the finger tapping and rest response blocks. By contrasting these blocks, we obtained the primary motor cortex areas activated by finger tapping.

### Extraction of betas (functional connectivity strengths) from IC-ACC

To evaluate potential negative correlations between the left DLPFC and the sgACC in the IC-ACC, we compared the personalized selection of targets with a selection of targets based on fixed MNI coordinates, as described in a recent study (Tik et al., 2017). We used MarsBar (Brett et al.,2017) to extract the parameter estimates (beta weights) of the left DLPFC locations at which we delivered the real rTMS stimulation, and of standard MNI locations of the DLPFC and sgACC (as described in Tik et al., 2017), after projecting the coordinates into the individual anatomical spaces of our participants. We used 2 mm radius ROI spheres to extract the betas from DLPFC regions. In Figure 3, note that personalized targets spread across the left DLPFC (individual, indDLPFC) and having a larger radius would have caused overlaps with the standard MNI DLPFC ROI (fixed, fxdDLPFC), diluting potential specific correlations with right sgACC (that we preserved with 5mm radius as in Tik et al., 2017 for comparison).

**Figure 3:**
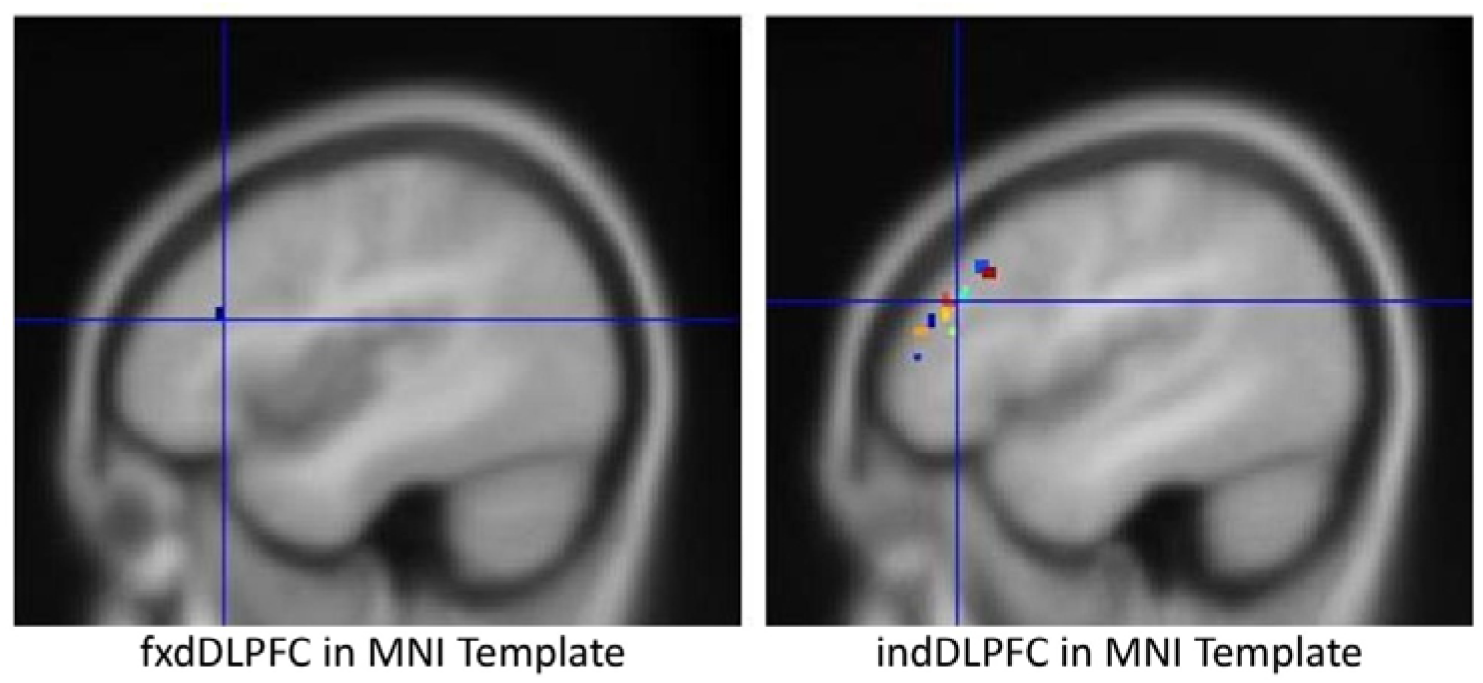
Figure showing the ROI of 2 mm radius around the standard MNI DLPFC coordinates in the left picture. On the right, are the 2 mm ROI around the individualized targets. Since selecting larger ROIs would have resulted in the overlap between the two for some cases, causing a dilution of results, we hence chose 2 mm ROI.

### Statistical Analysis

We used SPM12 to compare time windows of RS-fMRI across real and sham conditions, and only report results surviving a statistical threshold of whole brain p<0.05 FWE correction for multiple testing. We used SPSS and Matlab to run 2 sample t-tests to compare the scores from MADRS, HAM-D, YMRS, PANSS, PANAS, and VAS for real and sham stimulation sessions. Using MATLAB and R, we ran Pearson’s correlation tests between functional connectivity strengths of various brain regions.

## Results

Only 1 subject was excluded for not tolerating the stimulation, therefore data from 23 subjects were included in the final analysis. The mean age of the subjects was 25.8 ± 5.5 years with 9 female subjects (of 23 total).

### Target reproducibility

To establish the consistency of a method for personalized target selection, one has to first test whether or not target sites generated from different datasets of the same subject will coincide. To test the reproducibility of our target selection process, we repeated the steps of target selection on baseline RS-fMRI data from the day on which real stimulation was delivered. We then calculated the Euclidean distances between targets based on day 1 RS-fMRI data and those based on day 2 RS-fMRI data. The mean distance between the day 1 and day 2 targets was 10.9 mm, which was significantly less than 20 mm (p< 0.0001) as presupposed in the literature (see Discussion). Figure 4a presents a heat map of the distance between the targets for each subject. There are only three subjects for whom the distances between the day 1 and day 2 targets are slightly above 20 mm (diameter of the electric field sphere covered by figure 8 coils in Deng et al., 2013). Figure 4b presents a qualitative comparison of the modelled electric field (SimNIBS, Thielscher et al., 2015) generated by stimulation at targets for two example subjects.

**Figure 4a:**
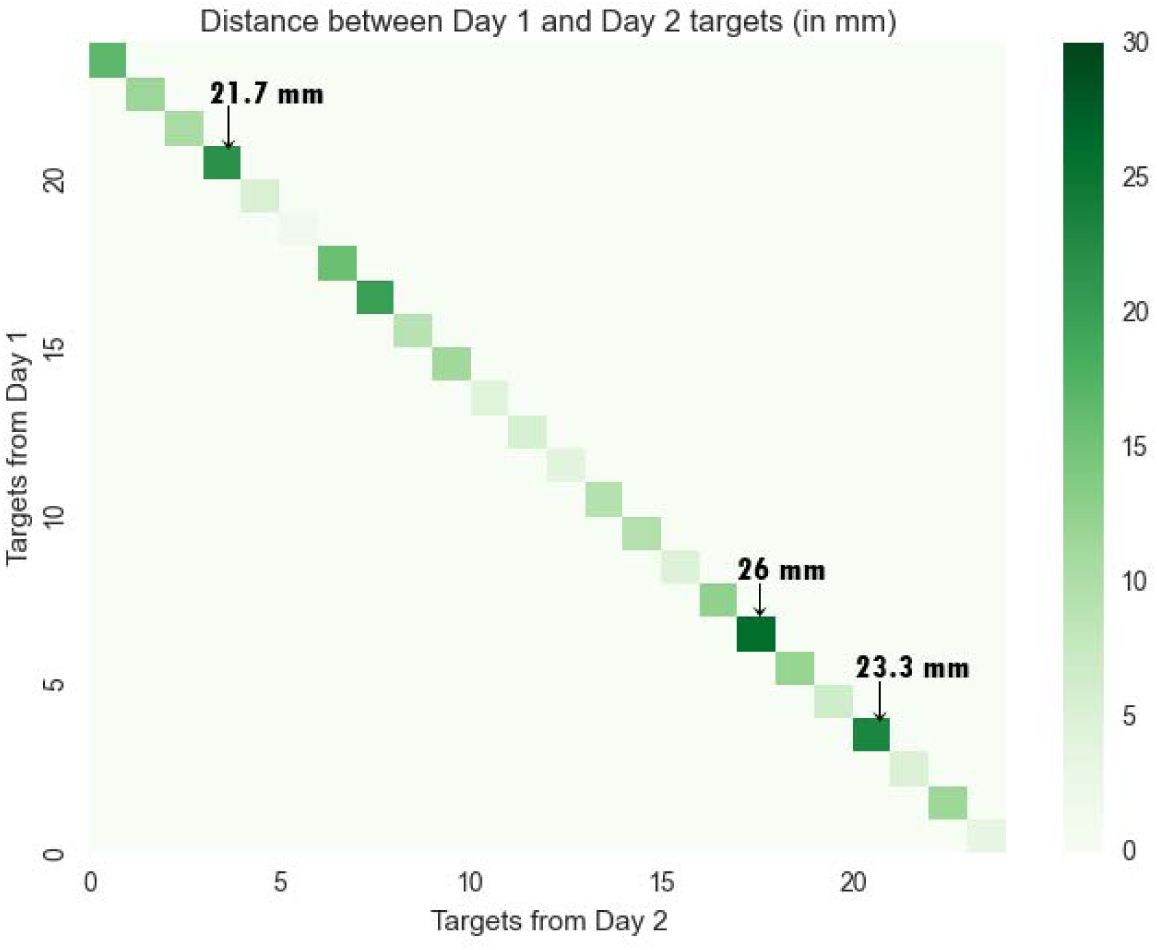
Heatmap of Euclidean distance between targets from day 1 RS-fMRI and targets from day 2 RS-fMRI. 3 subjects have a distance larger than 20 mm.

**Figure 4b:**
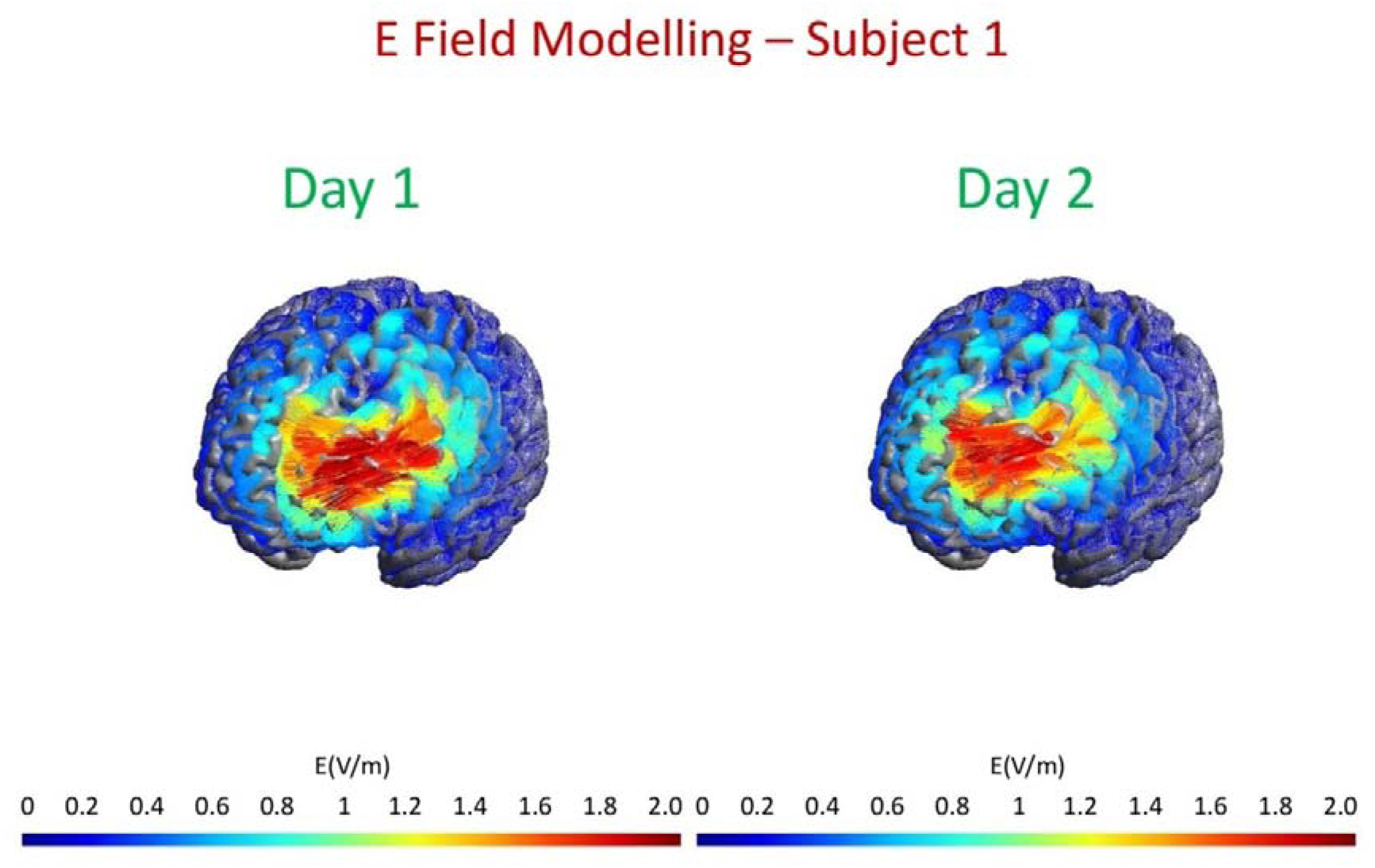
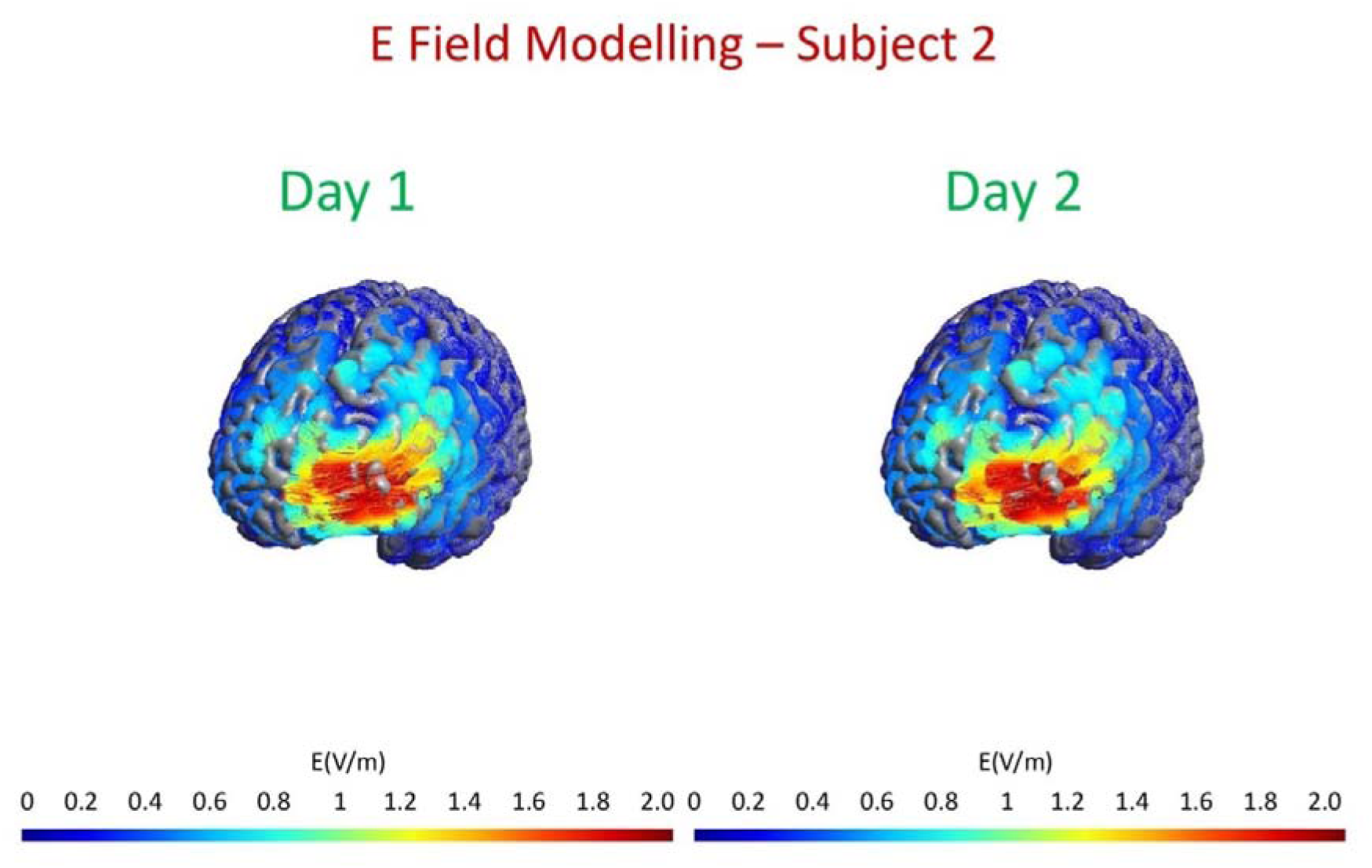
A qualitative description of electric field modeled (Thielscher et al., 2015) on the two targets from day 1 and day 2 RS-fMRI data. A quick looks reveals the similarity of the electric field when either of the targets are used.

### Target quality

The other important aspect of our target selection process is the quality of the target that it yields. Since our study involved healthy controls, it was not possible to obtain a clinical measure to determine the effectiveness of the target selection. However, as defined in Fox et al., 2012 & Weigand et al. 2017, effective targets for rTMS treatment of depression have higher negative functional connectivity to sgACC than less effective ones. Thus, we calculated the functional connectivity between the targets selected by our method (indDLPFC, see Methods) and the right sgACC, and also between fxdDLPFC and the sgACC. As expected, we report a higher negative connectivity between indDLPFC and sgACC compared to that between fxdDLPFC and sgACC (Figure 5), thus implying that indDLPFC would be more therapeutically effective compared to fxdDLPFC.

**Figure 5:**
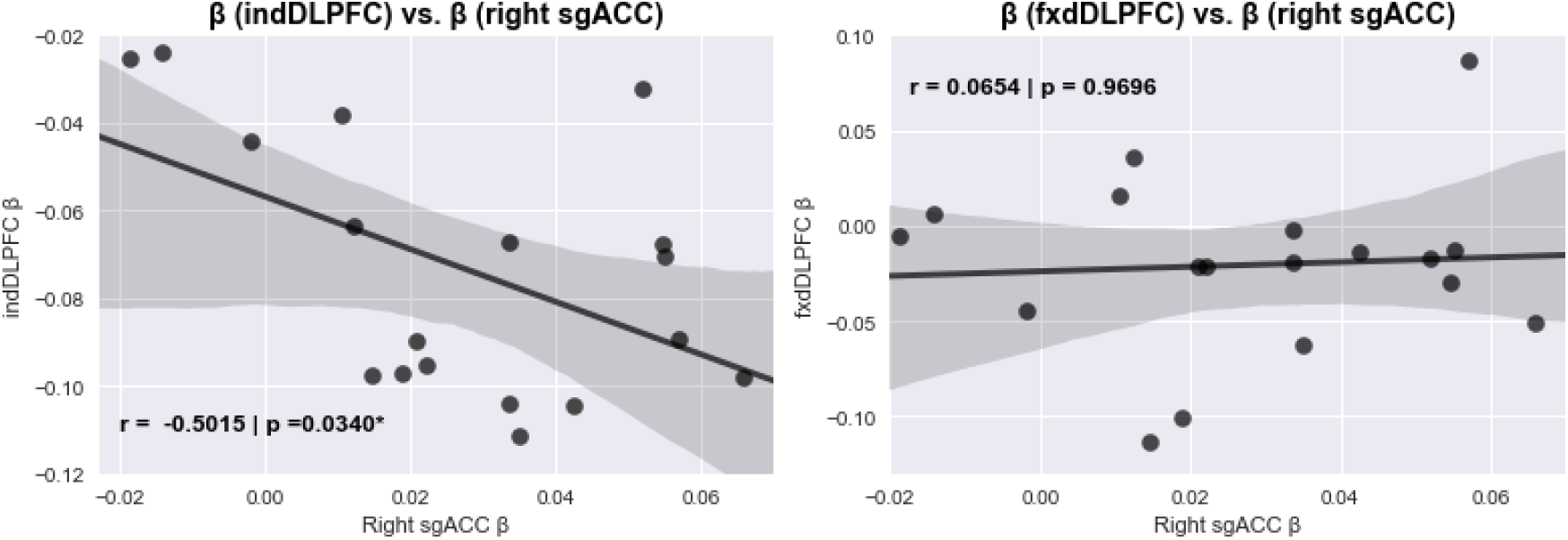
The plot on the left shows that targets identified using the described selection process have a much higher negative correlation with right sgACC (r = −0.5015, *p<0.05) than when the standard MNI left DLPFC coordinates are used (right plot). Since, antidepressant response of rTMS is linked to the connectivity of stimulated site and the sgACC; individualized targets would be promising for better therapeutic response than standard targets.

### Behavioural Scales Results

As expected, we did not see any differences in the MADRS, HAM-D, YMRS, PANSS, BDI II scales between day 2 and day 3 of stimulation.

However, the subjects reported a difference in the scalp discomfort experienced during real and sham stimulation (p<0.0001). Although they had different physical sensations arising from the two sessions, they expected both of the stimulation sessions to be equally effective (p>0.05, see Figure 6). This implies that our method of blinding successfully maintained similar levels of internal expectation even though they experienced a difference in scalp discomfort.

**Figure 6:**
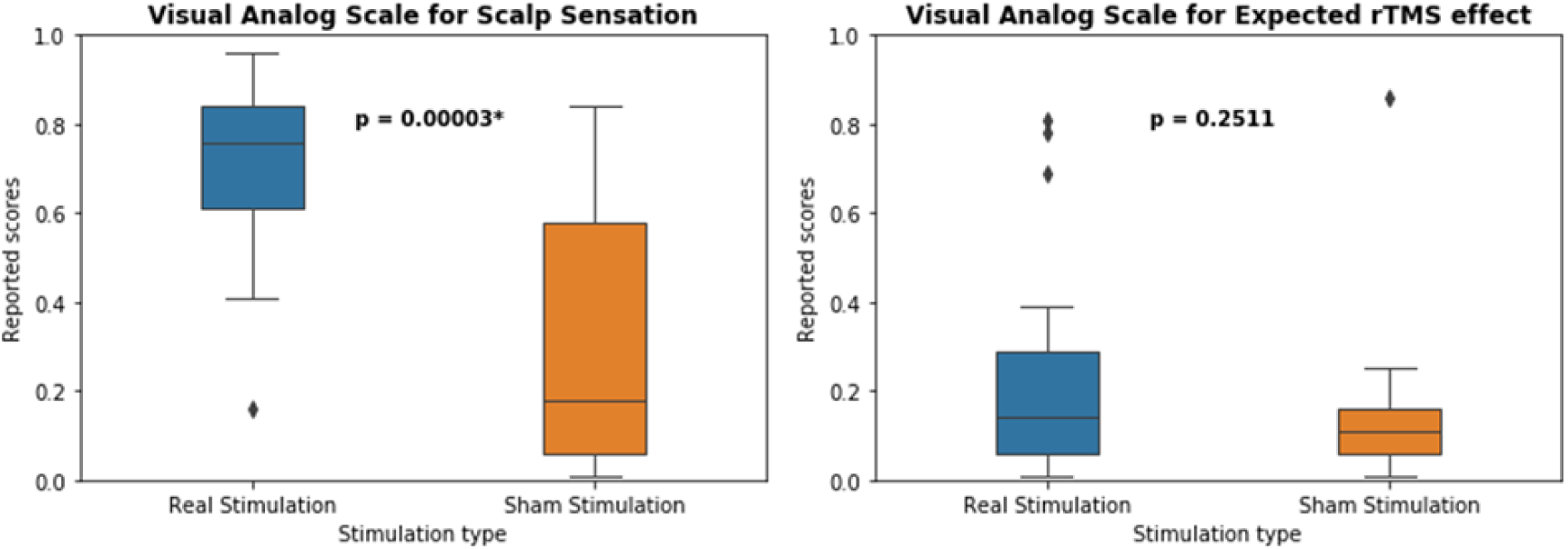
The plot on the left shows that the scalp discomfort experienced by the subjects was significantly higher during the real than sham condition. However, their internal expectation of the induced changes in their mental states did not differ across real and sham stimulation sessions.

### RS-fMRI functional connectivity changes post stimulation

To confirm the absence of false positive imaging results arising from nuisance movement, we compared the root mean square of framewise head displacement parameters (Van Dijk et al., 2012) from individual frames between the RS-fMRI sessions and found that the extent of motion did not differ across time windows or real and sham conditions.

To compare the changes in functional connectivity in the default mode network in real and sham conditions, we contrasted the four scanning sessions from real and sham stimulation for this network. We observed a robust decrease in functional connectivity specifically involving the sgACC (Figure 7a) and the ventral striatum (vStr) (Figure 7b) bilaterally during the R2-R1 contrast. This decrease in functional connectivity weakened over time and the connectivity in the ventral striatum returned to baseline level already in R3, while the decrease in the sgACC coupling persisted until R3, albeit less pronounced than during R2 (not shown).

**Figure 7:**
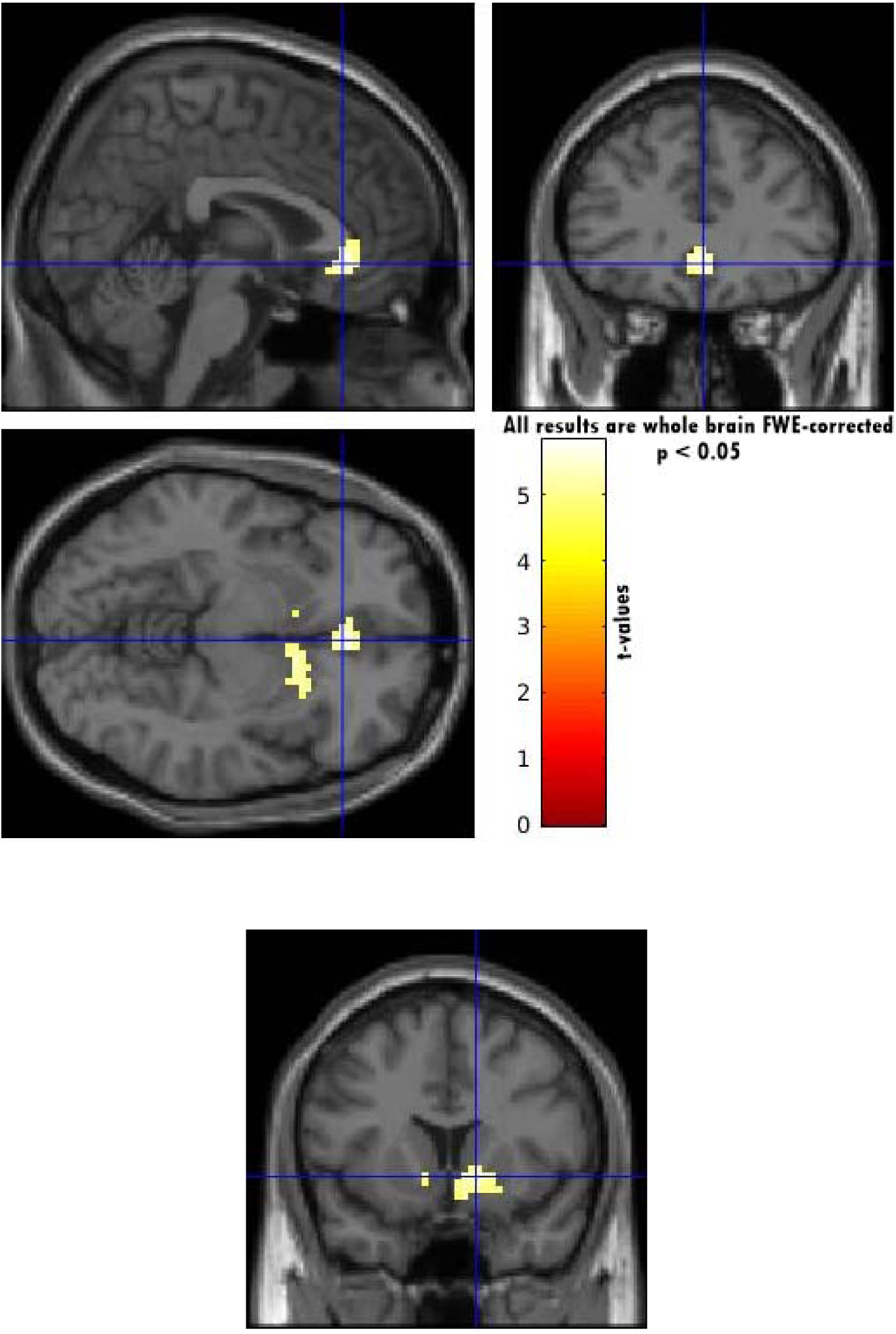
Bilateral sgACC (a) and vStr (b) regions, where functional connectivity of the default mode network decreases (real-sham contrast) during R2 RS-fMRI window (27-32 min) after rTMS stimulation. The color bar represents the t-values. All results shown are p<0.05 FWE corrected for whole brain; and are evidenced in real but not in sham condition.

### A predictor for rTMS response?

We were investigating healthy volunteers, and thus considered the dimensions of the TCI (novelty seeking, harm avoidance, reward dependence and persistence) informative. Therefore, we correlated the scores from this inventory with changes in functional connectivity strengths during R2 compared to R1. We identified a stronger negative correlation between harm avoidance (HA) and the decrease in functional connectivity strengths between the default mode network and the right sgACC (Figure 8) in the real condition (r = −0.5426, p value = 0.0164) but not in the sham condition (r = −0.2660, p value = 0.2710).

**Figure 8:**
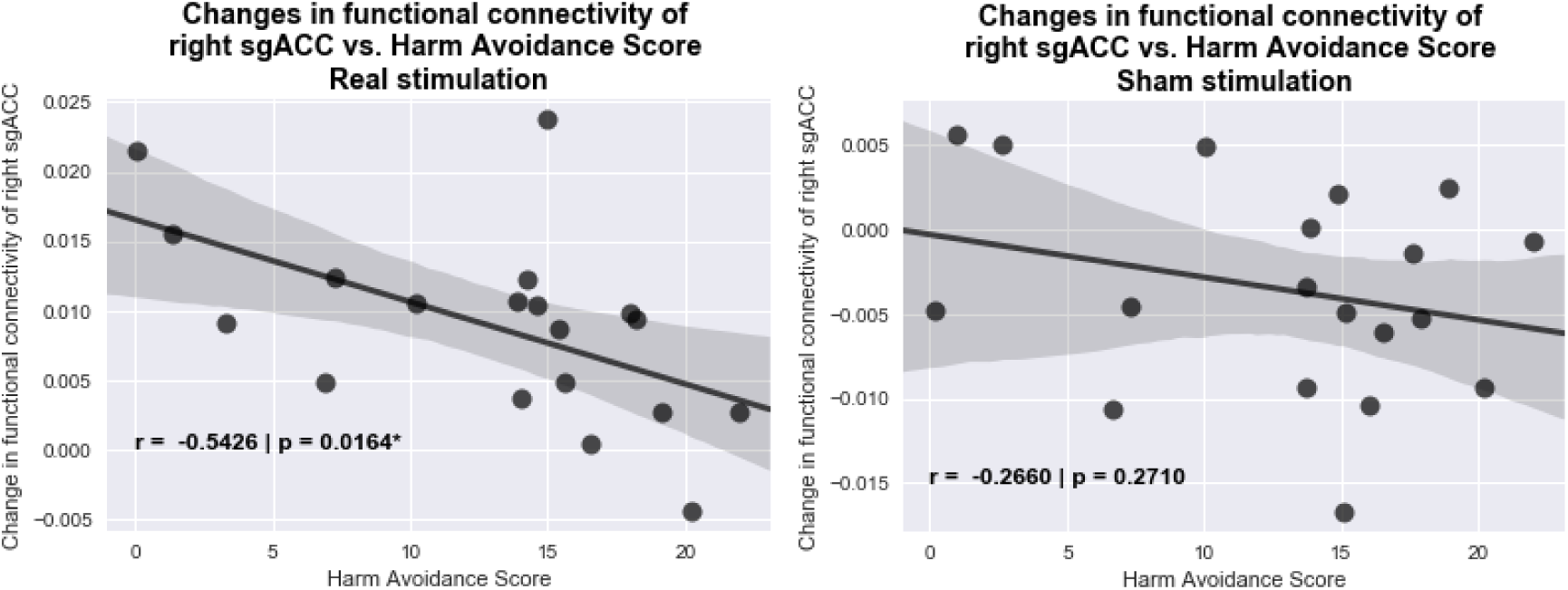
A negative correlation between harm avoidance score and the reduction in the functional connectivity of the default mode network at the subgenual ACC during R2 (27-32 min) from the R1 (10-15 min) RS-fMRI windows, indicates that the lower the subject scores on the harm avoidance scale, the higher was the uncoupling resulting from rTMS.

### Behavioural correlates of functional connectivity changes

The PANAS is a psychometric scale that measures both positive and negative affect. It is comprised of 20 items and includes two emotional dimensions: positive and negative. Ten items were used to evaluate negative affect and the remaining 10 items were used to evaluate positive affect. The raw score for each item ranged from one to five (Liu et al., 2012). We correlated the changes in scale scores (post stimulation – pre stimulation scores) with the functional connectivity of the default mode network in the right NAcc or the right sgACC extracted from beta weights. We report a trend of positive correlation (r = 0.3894, p value = 0.0993) of functional connectivity strengths in the right vStr during R2 with changes in negative affect scores, in the real condition but not in the sham condition (Figure 9).

**Figure 9:**
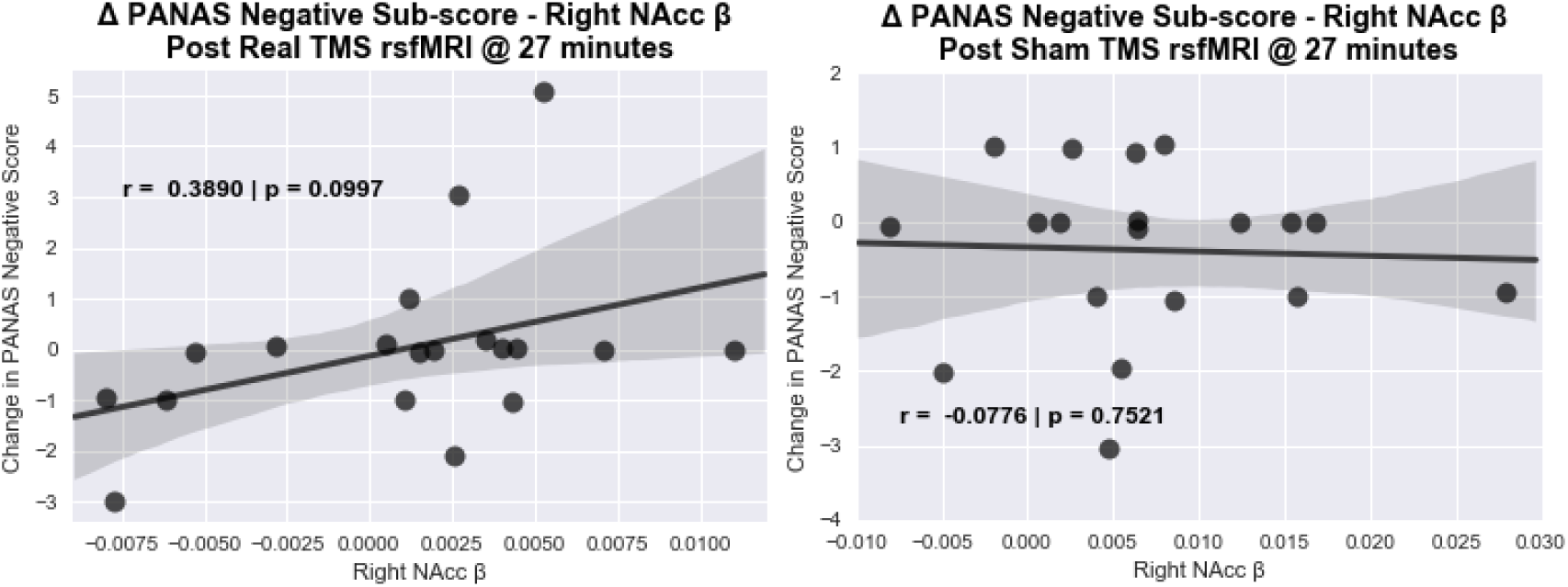
A trend of positive correlation observed between the functional connectivity strength of the default mode network at the right NAcc and the changes in negative affect score on PANAS scale occurs in real rTMS (left) but not in sham rTMS (right) condition. The correlation shows a transient effect during R2 RS-fMRI window (27-32 min) after rTMS: subjects who reported less negative emotions after stimulation showed less coupling between right NAcc and the default mode network.

## Discussion

In this study, we employed a novel personalized method to select the most promising left DLPFC spot that functionally relates to the sgACC. After verifying the necessary validation steps, we identify robust uncoupling of the default mode network and the subgenual ACC and ventral striatum in the real rTMS condition only. Considering the relevance of the sgACC for the pathophysiological changes underlying depression and antidepressant response, we investigated the strength of these effects in the personality dimension of HA. This dimension was selected considering its potential predictive value for rTMS response extending to healthy volunteers. As expected, subjects with higher HA scores presented less rTMS driven effects in the sgACC, which is in line with the literature investigating healthy and clinical samples (Abrams et al., 2004, Chen et al., 2015). Moreover, we identified a correlation trend of these transient effects in the ventral striatum and how subjects rated their negative emotions’ judgment.

For this novel target selection process in the left DLPFC to be reliable and of practical use, it is important that we can reproduce the target for an individual with an independent dataset i.e. RS-fMRI data from the same individual should yield similar targets. The targets should be analogous enough that stimulating at either of them produces comparable changes, when evidenced by electric field modelling. Previous research on modelling of electric fields (Deng et al., 2013) has shown that a figure 8 TMS coil covers electric field range that has a diameter of 20 mm (i.e. at 10 mm from the point of stimulation, the strength of the electric field is half maximum in all directions). Therefore, we used this distance as a reference for the reproducibility of our target effects. When calculating the Euclidean distances between targets resulting from two independent datasets, we have shown that they were significantly less than 20 mm apart from each other. This indicates that even if the target slightly shifted on the day of stimulation, it largely remained within the 20 mm electric field ambitioned by real rTMS. Thus, based on the above argument, our target selection is reproducible for practical purposes of personalized rTMS use.

Our new method employs individual subject’s RS-fMRI data at baseline to identify the strongest connectivity node in the left DLPFC anticorrelated to the sgACC. This guidance is of pivotal therapeutic significance in depression, as proposed by Fox and colleagues (Fox et al., 2012, Fox et al., 2013, Weigand et al., 2017). Weigand and colleagues have shown that for patients with depression, the higher the negative correlation between the stimulated left DLPFC region and the sgACC, the higher is the clinical improvement in response to rTMS. By doing so in healthy volunteers, we have shown that individualized targets selected by our method (indDLPFC) have a higher overall negative correlation with right sgACC (independent coordinates from Tik et al., 2017) than the fixed left DLPFC target (fxdDLPFC). Based on the results from Weigand et al. 2017, one could imply that determining indDLPFC targets for rTMS might result in more promising targets for clinical use.

We found a robust decrease of functional connectivity after real rTMS (R2) in the default mode network and the sgACC as well as the vStr. This decrease persists in the sgACC, albeit less pronounced, but in vStr it returns to normal during R3. We had a similar study design to Tik and colleagues (Tik et al., 2017), although our study protocol tracked the changes for a longer period of time - namely RS-fMRI windows of 10-15, 27-32, and 45-50 min - and investigated another resting state network. They reported an increase in the functional connectivity of the sgACC to a network consisting of the dorsal cingulate cortex, posterior dorsomedial prefrontal cortex, DLPFC, inferior parietal lobule, inferior frontal cortex and posterior temporal lobes. Moreover, they applied over a standard target in the left DLPFC about a third of the rTMS protocol that is clinically used to treat depression and at lower RMT (80%). This might at least partially explain the opposite direction of effects seen at the sgACC in comparison to our findings.

Several studies have observed higher sgACC functional connectivity in depression and its role in predicting antidepressant response (Drevets, 2000; Hamani et al., 2011; Dutta et al., 2014; Herringa et al., 2013; Gaffrey et al., 2012; Sheline et al., 2010, Greicius et al., 2007; Nugent et al., 2016; Jacon et al., 2016; Connolly et al., 2013; Musgrove et al., 2015; Baeken et al., 2015; Nugent et al., 2015). Further studies have shown abnormal sgACC connectivity with the default mode network during depression (Berman et al., 2011; Hamilton et al., 2011; Zhu et al., 2012). In this context, our results align well with the consequences of decoupling the default mode network and the sgACC after rTMS. Moreover, brain stimulation studies have reported a decreased activity in sgACC in response to deep brain stimulation (Berlim et al., 2014; Hamani et al., 2011). The NAcc, located at the vStr, is an integral component of the reward system and a recent study (Gong et al., 2016) has demonstrated disrupted reward circuit to be associated with depression severity activity of the NAcc region is related to anhedonia, a symptom of depression (Nauczyciel et al., 2013). A recent study also reported that its functional connectivity with stimulated left DLPFC predicts the antidepressant response of rTMS (Du et al., 2018). Given the importance of the sgACC and the NAcc regions in the pathophysiology of depression, the changes reported here in the sgACC and the vStr hold significance for the mechanisms underlying antidepressant effects. As both the sgACC and the NAcc are targets for deep brain stimulation (Mayberg et al., 2005; Lozano et al., 2008; Bewernick et al., 2012), our results demonstrate that it is possible to decrease the functional connectivity of these deep brain structures using non-invasive cortical rTMS.

It is important to note that our results are also in contrast to that reported by Taylor et al., 2018. Although they reported a decrease in functional connectivity of subgenual ACC, they found such changes in both sham and active conditions. We, however, found differences exclusively in the active stimulation condition. This discrepancy might stem from the fact that their results are from a patient population who were on a stable dosage of medication, whereas ours are from healthy volunteers under a single session of personalised rTMS. Our results in conjunction with those from Taylor and colleagues (Taylor et al., 2018) suggest that a decrease in functional connectivity of sgACC might be a shared mechanism implicated in antidepressant effects, both through pharmacological and rTMS interventions.

In light of the changes of functional connectivity of sgACC and vStr, we further considered the potential behavioural changes that could have resulted from rTMS stimulation. The negative correlation between the harm avoidance scores and the changes in functional connectivity in the real stimulation condition, but not in sham, indicate that harm avoidance score seems to predict the extent of the rTMS response in healthy subjects. This is an interesting result in light of a previous study (Mulder et al., 2006), which reported that harm avoidance was a negative predictor of response to antidepressant treatment, i.e. subjects with higher scores on harm avoidance responded poorly to antidepressant treatment. Apart from this, Ward et al., 2018 also reported a weak link between poor response to depression treatment and higher scores on neuroticism, which in turn positively correlates with harm avoidance scores (De Fruyt et al., 2000). Further studies by Quilty et al., 2008 and Seon-Young et al., 2007 also point towards a relation between high scores on neuroticism and low response to treatment of depression. Since our study is based on a cohort of healthy subject, we believe harm avoidance is a good proxy for neuroticism (given the positive correlation between the two parameters of personality). While we cannot, with certainty, claim that the harm avoidance scores can predict antidepressant response of rTMS, the correlation reported here warrants further studies to investigate the possibility of using the personality measure of harm avoidance as a predictor for antidepressant effects of rTMS.

Since the changes seen in the functional connectivity were on a time scale of minutes, we explored the changes in behaviour immediate to the stimulation. We hence explored the reported emotional state using PANAS scale, which we administered at the beginning and the end of the experiment. The PANAS is separated into negative and positive component scales, based on the literal meaning of the words, and asks the individual to rate their current experience of the emotion on a Likert scale from 1-5. For example, those words with a positive connotation (e.g. active, excited) are included in the positive scale score, where those with a negative connotation (e.g. upset, stressed) are included in the negative scale score. While the correlation between the negative subscale of PANAS and the functional connectivity strength of right vStr in R2 does not reach significance, there is a trend, which indicates that the lower the functional connectivity of the right vStr the more the subjects reported less negative emotions. This indicates that the changes in functional connectivity induced by the stimulation might be resulting in a less negative perception of emotions by the subjects and point towards a hypothesis that rTMS influences behaviour by causing changes in functional connectivity.

In this study, we developed a novel rTMS target selection process with the ultimate future goal of boosting the therapeutic efficiency of rTMS treatment for depression. However, since the study was initially performed with healthy subjects, it is not possible to comment on the clinical efficacy of stimulating at such targets in any way. Moreover, we cannot claim that the same functional connectivity differences seen in healthy participants will be sustained in the clinical sample. For that, we expect to validate and extend this method to patients in the near future. Another limiting aspect is that although with irrelevant current density, that did not resulted in functional connectivity differences, the sham condition was in fact an active sham condition. The use of a passive sham coil is unlikely to result in different results from those presented here, but this should nevertheless be rigorously tested. Lastly, the trend in correlation we identify in self-rated emotional judgement was not statistically significant. We consider two possible explanations for that, one being the sample size is limited in power, the other being that participants are healthy and only one session of rTMS is insufficient to exhibit stronger effects. However, new insights into the mechanisms associated with a full session of HF rTMS can be gained with our results.

## Conclusions

We present a new method of personalized target selection with utility for future rTMS studies. We have shown that the method is reproducible for practical purposes and allows targeting regions in left DLPFC that are most highly anticorrelated to sgACC. Although at present we cannot claim that this targeting process will be of higher therapeutic value based on our results, it withholds great promise of therapeutic use in patient cohorts to have therapeutic benefits compared to current methods of targeting left DLPFC. We also showed changes in the functional connectivity of the default mode network with the sgACC and the vStr, two brain regions implicated in the pathophysiology and treatment response of depression. Our study has shown that it is in fact possible to specifically influence the functional connectivity of both these deeper regions by stimulating non-invasively at cortical targets. And finally, we have shown that 1) the lower the harm avoidance score the stronger the rTMS effects in the sgACC 27-32 min after stimulation and 2) the lower the functional connectivity of the default mode network and the right NAcc 27-32 min after stimulation the less negative the subjects perceive their emotions, giving further insights on the mechanisms by which rTMS might be influencing behaviour.

## Acknowledgement

This work was supported by the German Federal Ministry of Education and Research (Bundesministerium fuer Bildung und Forschung, BMBF: 01 ZX 1507, “PreNeSt - e:Med”).

## Author Contributions

RGM, AA, and WP designed the study; AS, TEG, and GS collected and analyzed the data; AS wrote the manuscript and prepared the figures under the supervision of RGM. All authors reviewed the manuscript before submission.

